# nanoBERT: A deep learning model for gene agnostic navigation of the nanobody mutational space

**DOI:** 10.1101/2024.01.31.578143

**Authors:** Johannes Thorling Hadsund, Tadeusz Satława, Bartosz Janusz, Lu Shan, Li Zhou, Richard Röttger, Konrad Krawczyk

**Affiliations:** Department Mathematics and Computer Science, University of Southern Denmark; NaturalAntibody, Szczecin, Poland; Alector Therapeutics, 131 Oyster Point Blvd, Suite 600 South San Francisco, CA 94080

## Abstract

Nanobodies are a subclass of immunoglobulins, whose binding site consists of only one peptide chain, bestowing favorable biophysical properties. Recently, the first nanobody therapy was approved, paving the way for further clinical applications of this antibody format. Further development of nanobody-based therapeutics could be streamlined by computational methods. One of such methods is infilling - positional prediction of biologically feasible mutations in nanobodies. Being able to identify possible positional substitutions based on sequence context, facilitates functional design of such molecules. Here we present nanoBERT, a nanobody-specific transformer to predict amino acids in a given position in a query sequence. We demonstrate the need to develop such machine-learning based protocol as opposed to gene-specific positional statistics since appropriate genetic reference is not available. We benchmark nanoBERT with respect to human-based language models and ESM-2, demonstrating the benefit for domain-specific language models. We also demonstrate the benefit of employing nanobody-specific predictions for fine-tuning on experimentally measured thermostability dataset. We hope that nanoBERT will help engineers in a range of predictive tasks for designing therapeutic nanobodies.

**Availability:** https://huggingface.co/NaturalAntibody/

## 1 Introduction

Nanobodies (also called VHH, single domain antibodies) are a class of immunoglobulins, with binding sites consisting of only one polypeptide chain as opposed to two in canonical, human antibodies. The compact format bestows favorable biophysical properties, including high stability, greater tissue penetration and others [1,2]. Recent approval of the first nanobody therapeutic proved the feasibility of this format in human medicine [3], further reflected with an increasing patenting activity around nanobodies [4].

Design of novel therapeutic nanobodies can be facilitated by computational methods [5], such as structural modeling [6,7] or deimmunization/humanization [8,9]. Though cognate to antibodies, nanobodies bear a range of properties that set them apart from canonical antibodies [10–13]. For this reason, nanobodies can benefit from the breadth of computational approaches developed for antibodies, but require fine-tuning towards this specific molecule type [5,14].

Computationally guided antibody engineering requires a mutational map of the candidate molecule indicating feasible and non feasible mutations to help in activities such as humanization, affinity design, liabilities removal and others [8]. Such predictions are currently possible owing to the large number of Next-Generation Sequencing (NGS) samples deposited in the public domain, delineating the biologically acceptable mutations [15–17].

Mutational maps of antibodies can be created as position specific scoring matrices (PSSM) [18], or creating multiple sequence alignments (MSA) of millions of NGS sequences sharing a single germline origin [19,20]. Other studies have demonstrated the feasibility of gene agnostic language models of the human antibody space [21–24]. Here, large transformer models [25,26] are tasked with predicting obscured residues. Such transformer-based approaches have the advantage over PSSM/MSA based approaches in that they offer mutational predictions taking the entire context of the sequence rather than single positions.

The specific case of infilling nanobody sequences, could benefit from previous approaches developed for antibodies, either PSSM/MSA-based or transformer-based approaches. Though there exist reliable germline gene references for humans [27,28], the camelid, nanobody reference is not complete to the same extent [29]. In addition, the highly diverse germline segments, extended mutation hotspot regions and increased hypermutation frequency in VHH posed additional challenges for computational modeling [30]. Without gene assignment, creation of reliable PSSMs/MSAs is challenging. Therefore between PSSM/MSA and the transformer approach, only the latter remains as a feasible way of offering a mutational guide for nanobodies.

To address this issue, here we present a nanoBERT - a Bidirectional Encoder Representations from Transformers (BERT) model trained on ten million NGS sequences from the Integrated Nanobody Database for Immunoinformatics (INDI) [31]. Our model side-steps the need for reliable nanobody gene assignments offering a nanobody-specific mutational map of these molecules.

## 2 Materials and Methods

### Datasets employed

The training set for nanoBERT was compiled as the 10m nonredundant Next-generation sequencing nanobodies from INDI [31]. A total of 100,000 sequences were left as a validation set. We did not set aside a test set from this dataset as we believed that it would not be sound and instead opted from an entirely independent dataset (Table 1). Germline assignment of nanobody sequences were performed using ANARCI [32].

**Table 1.**
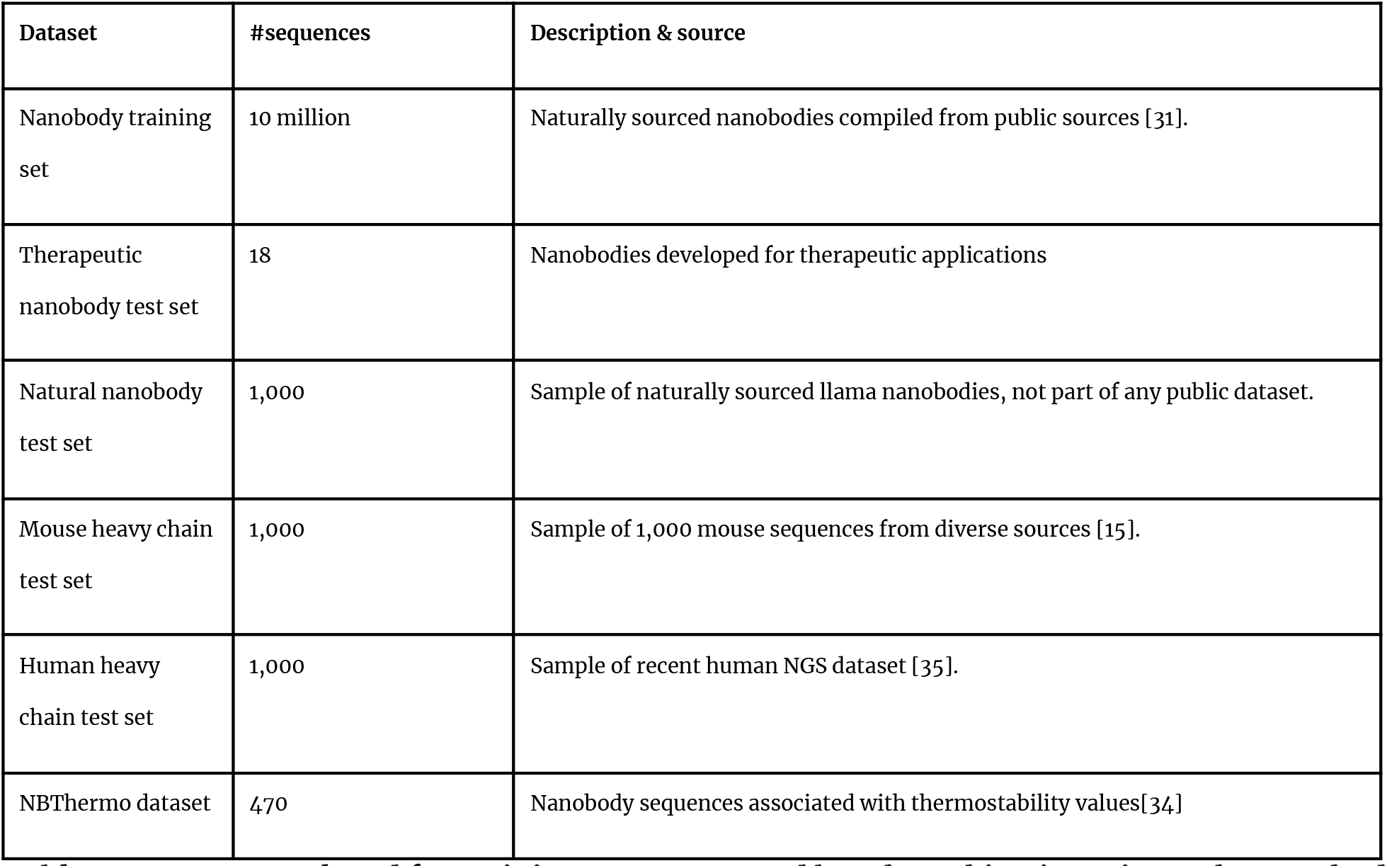
Datasets employed for training nanoBERT and benchmarking it against other methods. All sequences are non-redundant on the V-region level.

We used four test sets for infilling benchmarking (Table 1). As the blind test set we employed an internal NaturalAntibody dataset. The dataset comes from two llamas, whose repertoires were sequenced with approximately 500,000 sequences. This dataset does not form part of any public dataset and thus there should be no data leakage between test and train/validation. For computational expediency, a set of 1,000 nanobodies was sampled to constitute a natural test set. As the therapeutic test set we compiled a list of 18 therapeutic nanobodies from public sources [31,33]. These were Caplacizumab, Enristomig, Envafolimab, Gefurulimab, Gontivimab, Isecarosmab, Letolizumab, Lunsekimig, Ozekibart, Ozoralizumab, Porustobart, Rimteravimab, Sonelokimab, Tarperprumig and Vobarilizumab. Sonelokimab, Lunsekimig were multivalent, contributing three and two sequences respectively. This dataset was employed to indicate how well a model trained on natural sequences reconstructs therapeutic nanobody sequences. The mouse dataset was employed to contrast how well the models distinguish between different organisms through nativeness calculation. Here, we sampled a set of 1,000 non-redundant full V-region mouse sequences from OAS [15].

For the fine-tuning experimental dataset we employed the NBThermo dataset [34] which consists of nanobody sequences associated with thermostability measurements (Table 1).

### Transformer models employed

We trained a machine learning model based on the BERT paradigm following previous protocols, namely these of AntiBerta [21] and AntiBerty [23]. The objective task of our models was masked language modeling of 10m nanobody sequences from INDI (Table 1). We created two models, nanoBERT_big and nanoBERT_small. The nanoBERT_big model closely resembled the architecture of AntiBerta with 86m parameters and embedding size of 768. For comparison, we we also created nanoBERT_small, with 14m parameters and embedding size of 320 to check whether a smaller, more computationally efficient model would be comparable in terms of performance to the bigger one created using a standardized protocol.

For comparison of nanobody and human antibody models, we employed three language models trained solely on human data. We trained two heavy chain human models, human_320 and human_640, with 14m and 160m parameters respectively. The human_320 model resembles nanoBERT_small in terms of its architecture, but is trained on 25 million human heavy chains. The human_640 model is used to check whether scaling the number of parameters on the same dataset would achieve better prediction on nanobodies. As an external dataset, we also employed AbLang_heavy, which is a publicly available language model trained on human antibody sequences with the goal of sequence infiling. We note that there exist language models with inbuilt nanobody capacity, namely IgLM [24] and AbNativ [9], but we did not compare infilling against these two. The former performs sequence generation so it is not deterministic, making comparison unsound. To the best of our knowledge AbNativ provides a single sequence score indicating nativeness, without sequence infilling.

Finally, to compare the nanobody-specific language model to state-of the art protein language model not focused on a particular protein type, we employed ESM-2. We used the version with 650m parameters (*facebook/esm2_t33_650M_UR50D*) as the largest model we could compare head to head given our technical capacity.

### Fine-tuning experiments

We performed fine-tuning experiments on nativeness and thermostability calculations.

For the nativeness, we employed the test-set data from Table 1, to create two datasets, human vs nanobodies and mouse, as well as nanobodies vs human and mouse. Each dataset was then split in proportion of 8:1:1. Fine-tuning consisted of adding a four layer dense network with sigmoid output and binary cross-entropy loss.

In case of thermostability, we employed the NBThermo dataset (Table 1), splitting it between the Circular Dichroism, DSF (SYPRO) as well as putting all the data in one set. The thermostability values were mapped to range [0,1] and the same four layer dense network was employed for a regression task, but with linear output and mean squared error loss.

### Availability

We trained and benchmarked the models within the Hugging Face framework. We make our best nanobody model (nanoBERT_small) and a reference human heavy model (human_320) available under the link: https://huggingface.co/naturalantibody/. Model documentation contains a python notebook that can be easily run in Google Colab to demonstrate nanobody sequence infilling. The models can also be easily cloned from Hugging Face for custom applications such as fine-tuning, standardized within this framework.

## 3 Results

### Current camelid germlines are not sufficient to create reliable position specific scoring matrices

Before turning to language modeling we checked the possibility of mapping the nanobody space by clustering it into genes similarly to previous work on canonical antibodies (Młokosiewicz et al. 2022). Genes were assigned to the full INDI database using ANARCI [32]. Total of ∼94% of the database were correctly assigned to camelid genes, ∼6% as human and less than 0.01% as ‘cow’, ‘mouse’, ‘pig’, ‘rabbit’ and ‘rhesus’.

The chains that were correctly recognized as camelids, were unequally distributed between five V alleles (see Table 2). The very uneven assignment of germlines suggested poor gene identification, and we proceeded to evaluate the assignment by plotting the clonal distribution.

**Table 2.** Camelid V gene frequency of the INDI database assigned by ANARCI.

| V Gene | Frequency |
| --- | --- |
| IGHV3-3*01 | 68% |
| IGHV3S53*01 | 20% |
| IGHV3S1*01 | 10% |
| IGHV3-1*01 | 2% |
| IGHV4S1*01 | <1% |

We assumed that framework portions of the sequence should undergo the least somatic hypermutation, and thus should have the minimal distance from the assigned germline. As the framework portions we employed framework 2 (FW2) and 3 (FW3) for each chain as they are present in most sequences (framework 1 is sometimes truncated in NGS). Chains without data in FW2 and FW3 were excluded. For each allele we identified the most frequent concatenated FW2+3 sequence. Assuming proper gene assignment, the most frequent FW2+3 sequence from NGS would be identical to the germline and centroid of the chains clustered to the gene. We then ordered all other FW2+3 clones by their distance to the assigned germline sequence and plotted their frequency (see Figure 1). IGHV4S1^*^01 was omitted due to the low number of chains clustered to the gene (Table 2).

**Figure 1.**
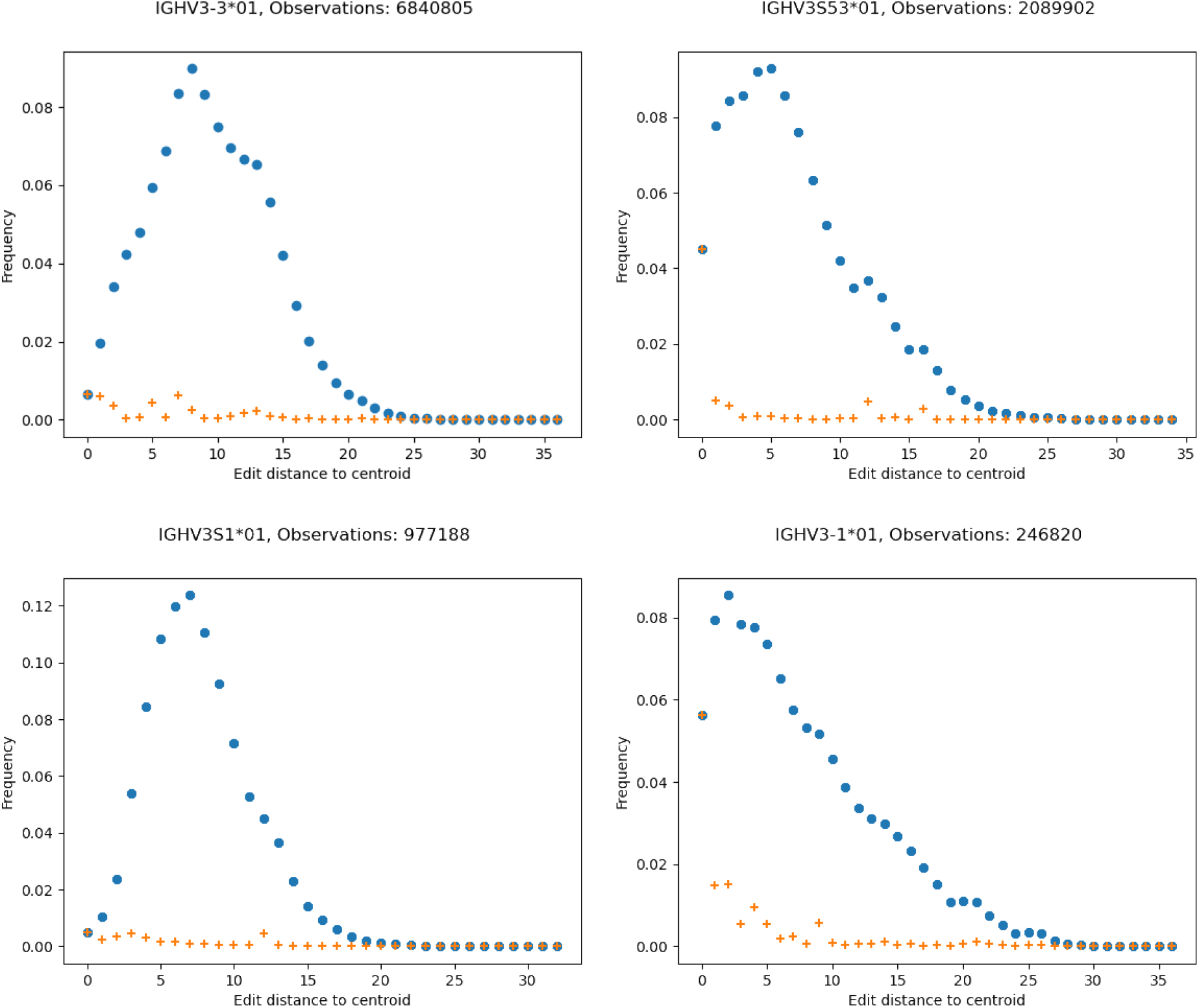
Frequency of nanobody sequences by mutational distance to the most frequent concatenated FW2+FW3 germline sequences. Blue circles indicate total frequency of all chains at a given distance to the central sequence, while red cross indicates the frequency of the most frequent unique sequence at each distance.

Assuming correct gene assignment, one would expect the germline FW2+FW3 being the centroid, and the slopes descending in Figure 1. The fact that there appear to be regions of higher density farther away from the germline, suggests genes/alleles that were not in our database. Therefore creating an infilling model using germline-based PSSM/MSA would be unsound in this case and should be tackled by using the NGS datasets to identify the novel germlines [36].

### Germline-agnostic transformer-based infilling of nanobody sequences

Since it was currently impossible to develop an infilling model employing the PSSM/MSA paradigm, we developed a transformer-based model. Specifically, we used BERT, which is a neural network architecture that enables representation of languages, and word prediction from both right and left context [25]. In nanoBERT each residue is considered as a word and each sequence as a sentence. nanoBERT was built using the same BERT architecture as AntiBerta [21] designed for human antibodies. We developed two models, nanoBERT_big (86m parameters) and nanoBERT_small (14m parameters) to check whether a more computationally efficient model performs on par with the larger one.

We benchmarked nanoBERT models against human-specific models and a protein-generalistic model on three datasets: natural nanobodies, natural human antibodies and therapeutic nanobodies (Table 3). Individual positions in each test-set sequence were obscured and the model was tasked with reconstructing them. If the top prediction matched the original sequence, it was counted as a match and mismatch otherwise. The accuracy for each region was calculated as the percentage of matches for each sequence divided by its length.

**Table 3:** Mean positional single amino acid prediction accuracy by IMGT-defined region. We count the accuracy of the query sequence matching the top prediction from a given language model.

| Natural llama nanobodies test set ( n = 1,000) |  |  |  |  |
| --- | --- | --- | --- | --- |
| Model | V region | FW | CDRs | CDR3 |
| nanoBERT_big | 76.6 | 88.0 | 44.2 | 31.7 |
| nanoBERT_small | 76.9 | 88.2 | 44.9 | 32.3 |
| human_320 | 64.7 | 76.5 | 31.3 | 19.6 |
| human_640 | 64.2 | 76.2 | 30.25 | 19.0 |
| AbLangHeavy | 64.4 | 76.4 | 30.3 | 18.8 |
| ESM-2 650M | 57.4 | 68.8 | 25.1 | 18.6 |
| Natural human antibodies test set (n=1,000) |  |  |  |  |
| Model | V region | FW | CDRs | CDR3 |
| nanoBERT_big | 64.5 | 73.7 | 37.7 | 25.9 |
| nanoBERT_small | 61.7 | 70.8 | 35.0 | 23.9 |
| human_320 | 91.4 | 97.2 | 74.8 | 57.3 |
| human_640 | 91.7 | 97.2 | 75.8 | 59.5 |
| AbLangHeavy | 91.7 | 97.0 | 76.4 | 60.8 |
| ESM-2 650M | 62.7 | 72.5 | 33.9 | 26.7 |
| Therapeutic nanobody test set (n=18) |  |  |  |  |
| Model | V region | FW | CDRs | CDR3 |
| nanoBERT_big | 77.4 | 88.2 | 46.0 | 30.2 |
| nanoBERT_small | 77.4 | 88.4 | 45.1 | 27.9 |
| human_320 | 73.9 | 86.4 | 37.1 | 23.1 |
| human_640 | 73.9 | 86.7 | 36.2 | 20.9 |
| AbLangHeavy | 70.1 | 82.1 | 35.0 | 23.0 |
| ESM-2 650M | 62.5 | 74.8 | 26.5 | 18.9 |

Performance on nanobodies dataset was supposed to contrast nanobody-specific models versus the nanobody-non-specific models. By symmetry, a test on human sequences was supposed to reflect whether human-specific models would outperform the nanobody-specific models. The therapeutic test set was supposed to indicate whether naturally-sourced predictions are useful in identifying mutations in nanobodies for therapeutic applications in humans.

On the natural nanobody dataset, which consisted of a sample of 1,000 sequences from sequencing of two llamas, the nanoBERT models outperform all the other models by a wide margin. The nanoBERT models achieve ca. 76% V region reconstruction versus ca. 64% for human models. The human antibody models perform better than ESM-2 (57.4% V-region accuracy), at nanobody infilling, reflecting the cognate nature of antibodies and nanobodies. The worst performance is noted for the CDR3 which is understandable as it is the most diverse region. Most notably, the small nanoBERT model appears to be firmly within the predictive range of the larger model, though it is more computationally efficient (thus we make available the nanoBERT_small model via Hugging Face).

Benchmarking the models on the human dataset, reverses the trend and now human antibody models perform much better than the nanobody-specific models. The human specific models achieve ca. 91% V region reconstruction accuracy versus 61-64% accuracy for nanoBERT models. The accuracy of nanoBERT models for the entire V-region is within the range of ESM-2 (62.5%). Therefore, the reflective case of nanobody models performing better on cognate antibodies than a generalistic protein model is not the case. Of note, the human-specific models achieve much better reconstruction rate on human sequences (ca. 91%) than nanoBERT on nanobodies (ca. 76%), which could be due to larger, more diverse datasets being used to train these.

Finally, we tested all models on infilling the 18 therapeutic nanobody sequences. Here, the performance gap between nanoBERT and human-specific models is much smaller than on natural datasets. The nanoBERT models achieve better prediction on the entire V-region (ca. 77% for nanoBERT models versus ca. 70-73% for human-specific models). The biggest performance gap is noted for the entire CDR regions with nanoBERT models achieving ca. 45% accuracy versus 35% accuracy for human-specific models. On the therapeutic set, all the models outperformed ESM-2 by a wide margin.

Therefore, these results demonstrate that nanobody-specific transformer models provide benefit in objective tasks of sequence infilling that could have an application in engineering such molecules.

### Single domain-based therapeutics employ nanobody hallmark residues

Application of infilling models to therapeutic nanobodies is an important problem, with potential for sequence liability removal within biologically relevant space. Cognate application is humanization - making the molecule resemble human amino acid distribution at certain positions [8]. Simply grafting the nanobody CDRs onto human frameworks is not efficient as binding specificity can be lost. There is much evidence and data collected on murine-based deimmunization where strategically placed substitutions maximize human nativeness, without compromising stability of the molecule [37]. However with only eighteen nanobody therapeutics, comparable knowledge on these molecules is still being developed.

To check to what extent existing therapeutic nanobody molecules reflect their natural versus human amino acid distribution, we plotted multiple sequence alignments of the therapeutic nanobodies and the closest human germlines (Figure 2). In all cases, the most mismatches with the human germline are accumulated in the framework 2 regions. Eight out of eighteen nanobody therapeutics have framework 2 hallmark motif FERF, as opposed to the typical human VGLW. Therefore current engineering choices take nanobody-preferred residues [38], even though these are not preferred substitutions according to the human amino acid distribution.

**Figure 2.**
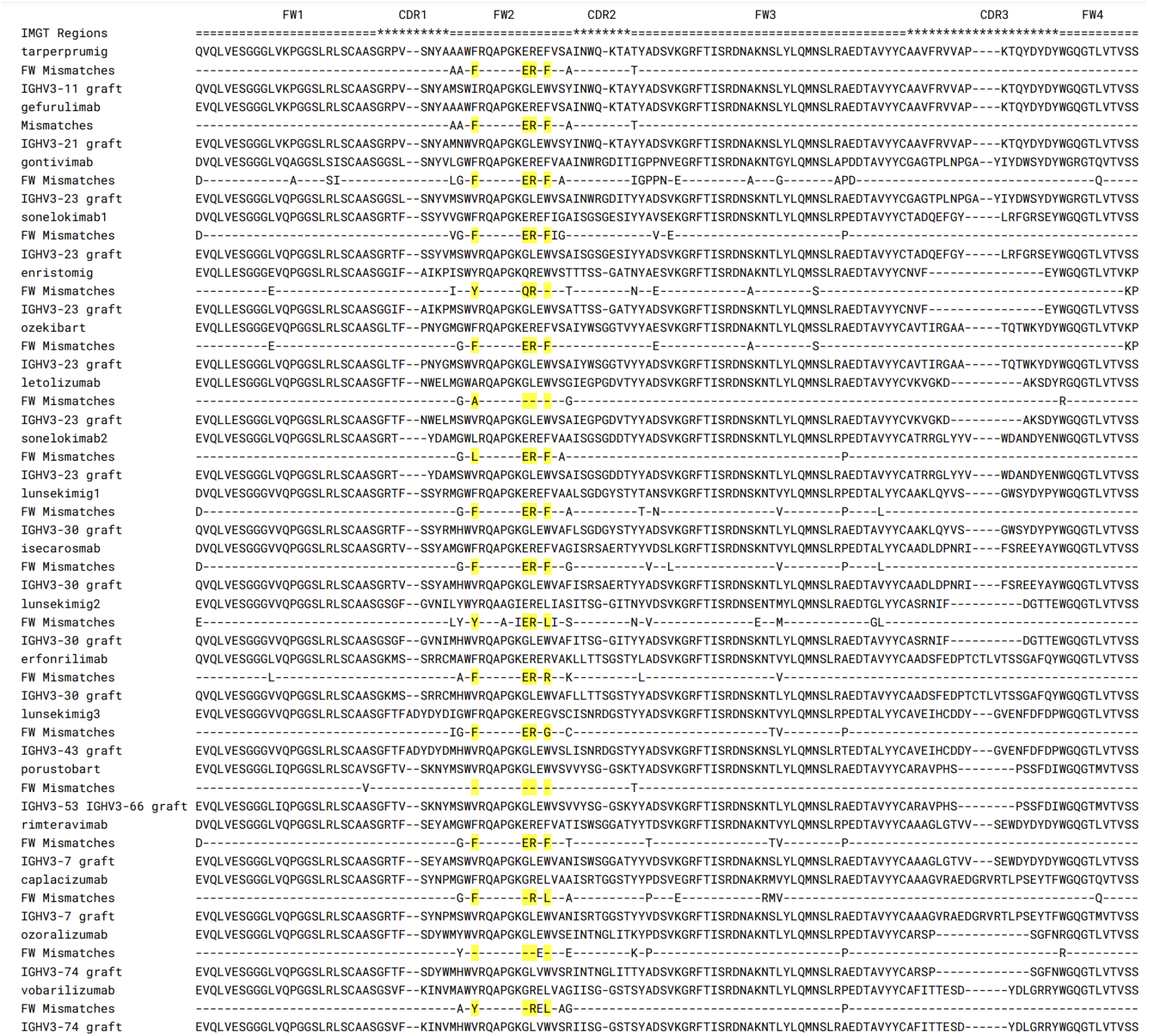
Multiple sequence alignment of nanobody-based therapeutics. For each nanobody-based therapeutic the closest human germline was identified. Graft of the IMGT CDRs is shown along with mismatches between the closest human framework and the original therapeutic. Nanobody hallmark residues are annotated in yellow.

Employing a nanobody-specific infilling model can better reflect amino acid distribution preferred to such residues. Contrasting such distributions with human variable regions [8] or human transformers (e.g. human_320) might provide the initial data basis on humanization of these molecules whilst clinical trial data on anti-drug antibodies to nanobodies is accumulated.

### Downstream zero shot and fine tuning prediction tasks

Self-supervised models can be applied to a range of problems in zero-shot fashion, predicting properties they were not specifically trained on [39]. Here we focused on nativeness, which is correlating aggregate positional residue predictions with attribution of sequences to a given species [40].

We calculate the nativeness as the sum of inverse exponents of the last layers in our nanoBERT_small and human_320 transformers. The lower the nativeness value, the closer the query sequence to the sequence distribution the model was trained on. We contrasted it with the single nanobody-specific nativeness score available, namely AbNativ [9]. Here nativeness is calculated using variation of the variational autoencoder, using either human or nanobody-based distributions. We contrast the nativeness for nanoBERT_small, human_320, AbNativ human and AbNativ nanobody, on our human, nanobody and mouse test sets (Figure 3).

**Figure 3.**
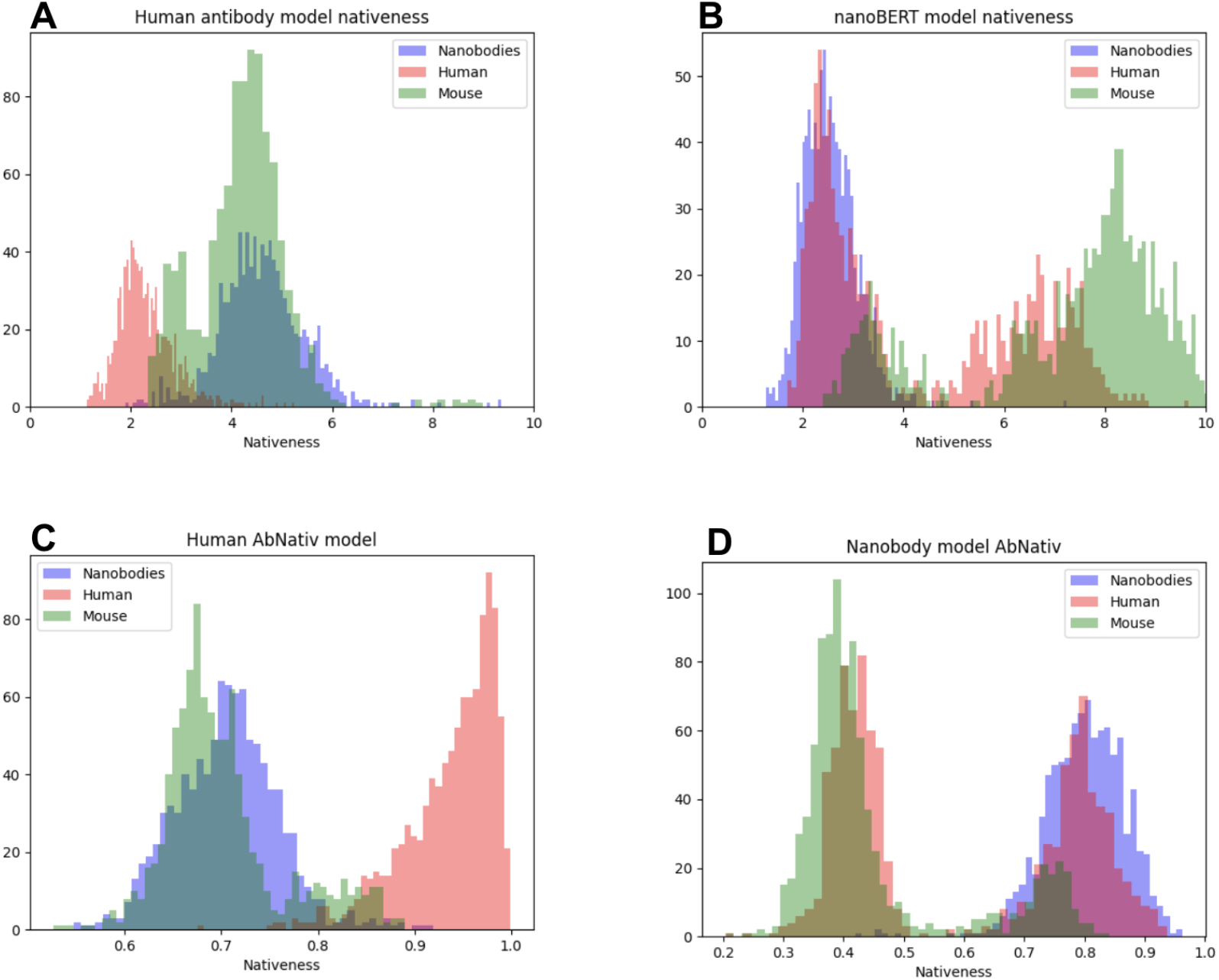
Nativeness calculation based on nanobody and human models. Nativeness was calculated using our nanoBERT_small and human_320 models and contrasted to AbNativ VHH (nanobody) and VH (antibody) models. Please note that the nativeness scales are not comparable between AbNativ and our models. In case of AbNativ, higher values are more native whereas in case of nanoBERT_small and human_320, lower values indicate higher nativeness. **A**. Nativeness calculated using the human_320. **B**. Nativeness calculated using nanoBERT_small **C**. Nativeness calculated using AbNativ VH score. **D**. Nativness calculated using AbNativ VHH score.

It is clear that the human models do a much better job in distinguishing human sequences from non-human ones (Figure 3A, 3C). By contrast, both nanobody models do quite poorly on zero-shot separation of our llama nanobodies from human or mice ones (Figure 3B, 3D). In both cases the nanobody distribution is fully mixed with the human one. It cannot be the case that there exist human sequences that are too similar to nanobody ones to be distinguished, otherwise the human-only models would likewise fail at separation.

We further checked the difficulty of the sequence separation problem in a fine-tuning fashion, by training a four layer dense head on top of either human_320 or nanoBERT_small. In each case, even training on a very small dataset (splitting our test sets in 8:1:1 fashion into human vs mouse and nanobody or nanobody vs human and mouse), yielded PROC AUCs approaching 1.0.

Species separation is a trivial problem that can be solved by non-machine learning methods using sequence alignment to closest germlines (Table 2). To test out transformers on a more challenging downstream task, we fine-tuned a predictor on the NBThermo dataset, that is a compilation of nanobody sequences associated with their melting temperatures [34]. The entire dataset we employed consisted of 470 sequences, with the experimental measurements of Circular Dichroism and DSF (SYPRO) having the most sequences, 263 and 240 respectively.

We split the fine-tuning into three cases. First, training and testing on Circular Dichroism, second, training and testing on DSF (SYPRO) and finally, training and testing on all the measurements. Sequences were split into train and test sets in 8:1:1 fashion and 90% sequence identity restriction was imposed. The melting temperatures were mapped to the range [0,1] and predicting the value in this range was the objective task of the head built on top of the transformer.

The models were applied to three splits of the NbThermo dataset. In each case, the model was fine-tuned five times and the Pearson correlation calculated on the test dataset. Multiple training sessions were applied to check the stability of the training given such a small dataset. As a comparison on the random performance on a given split, we also calculated the random baseline, sampling the prediction scores from a uniform random distribution in the range of [0,1]. To show the effect of pre-training, we also trained the same model architecture but without earlier pre-training. The results of the fine-tuning experiments are given in Table 4. Both the nanobody and human models outperform the random baseline and the model without pre-training. The nanobody-specific transformer appears to have better performance on the Circular Dichroism dataset, DSF (SYPRO) and has marginally worse performance on the all dataset. The human transformer achieves very poor performance on the DSF (SYPRO) dataset. Altogether the results suggest that there might be some benefit in employing a nanobody-specific model to fine-tune on nanobody-specific tasks. Nevertheless, it is still to be verified on a larger dataset how generalizable such models are actually.

**Table 4.** Fine tuning nanoBERT_small and human_320 on NbThermo dataset. ‘Only head’ column, indicates the performance of a model where no pre-training was performed. Random column indicates performance on the test set when a random number in range [0,1] was generated. Fine-tuning was performed five times on each model and each dataset and the mean value is reported.

| NbThermo subset | Pearson correlation |  |  |  |
| --- | --- | --- | --- | --- |
|  | nanoBERT_small | human_320 | Only Head | Random |
| Circular Dichroism (n=263) | $\mu = 0.5 (\sigma = 0.04)$ | $\mu = 0.39 (\sigma = 0.05)$ | $\mu = 0.06 (\sigma = 0.29)$ | $\mu = 0.062 (\sigma = 0.04)$ |
| DSF (SYPRO) (n=240) | $\mu = 0.41 (\sigma = 0.06)$ | $\mu = 0.07 (\sigma = 0.06)$ | $\mu = -0.02 (\sigma = 0.05)$ | $\mu = -0.194 (\sigma = 0.22)$ |
| All (n=470) | $\mu = 0.43 (\sigma = 0.03)$ | $\mu = 0.44 (\sigma = 0.03)$ | $\mu = 0.23 (\sigma = 0.10)$ | $\mu = 0.029 (\sigma = 0.11)$ |

## 4 Discussion

We demonstrated that a nanobody-specific transformer outperforms those trained on generalistic protein data, and those focused on human heavy chains. Such infilling models could be instrumental in providing mutational choices in the process of engineering these molecules.

The clearest application of infilling is with humanizing nanobodies, that is already being addressed by experimental [38] and machine learning protocols [8]. Choosing substitutions so as to best reflect human distribution, whilst maintaining the single-domain character of a nanobody therapeutic could be challenging. Tapping onto a model of human and nanobody distribution could indicate positions that are strongly preferred in one or the other, offering evidence in favor or against certain substitutions. Any benefits of such data-driven approaches will be properly benchmarked when more immunogenicity data on nanobodies becomes available with future clinical trials.

Beyond infilling, applying models self-supervised on large sequence datasets, holds much potential in focusing these on much smaller experimental datasets. Though we have demonstrated benefits of using nanobody-specific datasets for fine-tuning, a much wider benchmark on encompassing multiple nanobody-specific experimental datasets would be necessary.

We make our human and nanobody models available via Hugging Face in hope that they will facilitate multiple therapeutic applications, both within the scope of sequence infilling and fine-tuning.

## References

1. Bannas P, Hambach J, Koch-Nolte F. Nanobodies and Nanobody-Based Human Heavy Chain Antibodies As Antitumor Therapeutics. Front Immunol. 2017;8: 1603.

2. Flajnik MF, Deschacht N, Muyldermans S. A case of convergence: why did a simple alternative to canonical antibodies arise in sharks and camels? PLoS Biol. 2011;9: e1001120.

3. Morrison C. Nanobody approval gives domain antibodies a boost. Nat Rev Drug Discov. 2019;18: 485–487.

4. Krawczyk K, Buchanan A, Marcatili P. Data mining patented antibody sequences. MAbs. 2021;13: 1892366.

5. Wilman W, Wróbel S, Bielska W, Deszynski P, Dudzic P, Jaszczyszyn I, et al. Machine-designed biotherapeutics: opportunities, feasibility and advantages of deep learning in computational antibody discovery. Brief Bioinform. 2022;23. doi:10.1093/bib/bbac267

6. Cohen T, Halfon M, Schneidman-Duhovny D. NanoNet: Rapid end-to-end nanobody modeling by deep learning at sub angstrom resolution. bioRxiv. 2021. p. 2021.08.03.454917. doi:10.1101/2021.08.03.454917

7. Abanades B, Wong WK, Boyles F, Georges G, Bujotzek A, Deane CM. ImmuneBuilder: Deep-Learning models for predicting the structures of immune proteins. Commun Biol. 2023;6: 575.

8. Sang Z, Xiang Y, Bahar I, Shi Y. Llamanade: An open-source computational pipeline for robust nanobody humanization. Structure. 2021. doi:10.1016/j.str.2021.11.006

9. Ramon A, Saturnino A, Didi K, Greenig M, Sormanni P. AbNatiV: VQ-VAE-based assessment of antibody and nanobody nativeness for engineering, selection, and computational design. bioRxiv. 2023. p. 2023.04.28.538712. doi:10.1101/2023.04.28.538712

10. Mitchell LS, Colwell LJ. Analysis of nanobody paratopes reveals greater diversity than classical antibodies. Protein Eng Des Sel. 2018;31: 267–275.

11. Mitchell LS, Colwell LJ. Comparative analysis of nanobody sequence and structure data. Proteins. 2018;86: 697–706.

12. Gordon GL, Capel HL, Guloglu B, Richardson E, Stafford RL, Deane CM. A comparison of the binding sites of antibodies and single-domain antibodies. Front Immunol. 2023;14: 1231623.

13. Li X, Duan X, Yang K, Zhang W, Zhang C, Fu L, et al. Comparative Analysis of Immune Repertoires between Bactrian Camel’s Conventional and Heavy-Chain Antibodies. PLoS One. 2016;11: e0161801.

14. Norman RA, Ambrosetti F, Bonvin AMJJ, Colwell LJ, Kelm S, Kumar S, et al. Computational approaches to therapeutic antibody design: established methods and emerging trends. Brief Bioinform. 2020;21: 1549–1567.

15. Kovaltsuk A, Leem J, Kelm S, Snowden J, Deane CM, Krawczyk K. Observed Antibody Space: A Resource for Data Mining Next-Generation Sequencing of Antibody Repertoires. J Immunol. 2018;201: 2502–2509.

16. Olsen TH, Boyles F, Deane CM. Observed Antibody Space: A diverse database of cleaned, annotated, and translated unpaired and paired antibody sequences. Protein Sci. 2022;31: 141–146.

17. Briney B. AntiRef: reference clusters of human antibody sequences. Bioinformatics Advances. 2023; vbad109.

18. Smith MD, Case MA, Makowski EK, Tessier PM. Position-Specific Enrichment Ratio Matrix scores predict antibody variant properties from deep sequencing data. Bioinformatics. 2023;39. doi:10.1093/bioinformatics/btad446

19. Młokosiewicz J, Deszyński P, Wilman W, Jaszczyszyn I, Ganesan R, Kovaltsuk A, et al. AbDiver-A tool to explore the natural antibody landscape to aid therapeutic design. Bioinformatics. 2022. doi:10.1093/bioinformatics/btac151

20. Schmitz S, Soto C, Crowe JE Jr, Meiler J. Human-likeness of antibody biologics determined by back-translation and comparison with large antibody variable gene repertoires. MAbs. 2020;12: 1758291.

21. Leem J, Mitchell LS, Farmery JHR, Barton J, Galson JD. Deciphering the language of antibodies using self-supervised learning. Patterns (N Y). 2022;3: 100513.

22. Olsen TH, Moal IH, Deane CM. AbLang: an antibody language model for completing antibody sequences. Bioinformatics Advances. 2022;2: vbac046.

23. Ruffolo JA, Gray JJ, Sulam J. Deciphering antibody affinity maturation with language models and weakly supervised learning. arXiv [q-bio.BM]. 2021. Available: http://arxiv.org/abs/2112.07782

24. Shuai RW, Ruffolo JA, Gray JJ. Generative Language Modeling for Antibody Design. bioRxiv. 2021. p. 2021.12.13.472419. doi:10.1101/2021.12.13.472419

25. Devlin J, Chang M-W, Lee K, Toutanova K. BERT: Pre-training of deep bidirectional Transformers for language understanding. 2018. doi:10.48550/ARXIV.1810.04805

26. Vaswani A, Shazeer N, Parmar N, Uszkoreit J, Jones L, Gomez AN, et al. Attention is all you need. Adv Neural Inf Process Syst. 2017;30. Available: https://proceedings.neurips.cc/paper/7181-attention-is-all-you-need

27. Smakaj E, Babrak L, Ohlin M, Shugay M, Briney B, Tosoni D, et al. Benchmarking immunoinformatic tools for the analysis of antibody repertoire sequences. Bioinformatics. 2020;36: 1731–1739.

28. Lefranc MP, Giudicelli V, Ginestoux C, Bodmer J, Müller W, Bontrop R, et al. IMGT, the international ImMunoGeneTics database. Nucleic Acids Res. 1999;27: 209–212.

29. Tu Z, Huang X, Fu J, Hu N, Zheng W, Li Y, et al. Landscape of variable domain of heavy-chain-only antibody repertoire from alpaca. Immunology. 2020;161: 53–65.

30. Nguyen VK, Hamers R, Wyns L, Muyldermans S. Camel heavy-chain antibodies: diverse germline V(H)H and specific mechanisms enlarge the antigen-binding repertoire. EMBO J. 2000;19:921–930.

31. Deszyński P, Młokosiewicz J, Volanakis A, Jaszczyszyn I, Castellana N, Bonissone S, et al. INDI-integrated nanobody database for immunoinformatics. Nucleic Acids Res. 2022;50: D1273–D1281.

32. Dunbar J, Deane CM. ANARCI: antigen receptor numbering and receptor classification. Bioinformatics. 2016;32: 298–300.

33. Raybould MIJ, Marks C, Lewis AP, Shi J, Bujotzek A, Taddese B, et al. Thera-SAbDab: the Therapeutic Structural Antibody Database. Nucleic Acids Res. 2020;48: D383–D388.

34. Valdés-Tresanco MS, Valdés-Tresanco ME, Molina-Abad E, Moreno E. NbThermo: a new thermostability database for nanobodies. Database. 2023;2023. doi:10.1093/database/baad021

35. Jaffe DB, Shahi P, Adams BA, Chrisman AM, Finnegan PM, Raman N, et al. Functional antibodies exhibit light chain coherence. Nature. 2022;611: 352–357.

36. Ralph DK, Matsen FA 4th. Per-sample immunoglobulin germline inference from B cell receptor deep sequencing data. PLoS Comput Biol. 2019;15: e1007133.

37. Tennenhouse A, Khmelnitsky L, Khalaila R, Yeshaya N, Noronha A, Lindzen M, et al. Computational optimization of antibody humanness and stability by systematic energy-based ranking. Nat Biomed Eng. 2023. doi:10.1038/s41551-023-01079-1

38. Saerens D, Pellis M, Loris R, Pardon E, Dumoulin M, Matagne A, et al. Identification of a universal VHH framework to graft non-canonical antigen-binding loops of camel single-domain antibodies. J Mol Biol. 2005;352: 597–607.

39. Meier J, Rao R, Verkuil R, Liu J. Language models enable zero-shot prediction of the effects of mutations on protein function. Adv Neural Inf Process Syst. 2021. Available: https://proceedings.neurips.cc/paper_files/paper/2021/hash/f51338d736f95dd42427296047067694-Abstract.html

40. Wollacott AM, Xue C, Qin Q, Hua J, Bohnuud T, Viswanathan K, et al. Quantifying the nativeness of antibody sequences using long short-term memory networks. Protein Eng Des Sel. 2019;32: 347–354.

